# Denoising, Deblurring, and optical Deconvolution for cryo-ET and light microscopy with a physics-informed deep neural network DeBCR

**DOI:** 10.1101/2024.07.12.603278

**Authors:** Rui Li, Artsemi Yushkevich, Xiaofeng Chu, Mikhail Kudryashev, Artur Yakimovich

## Abstract

Computational image-quality enhancement for microscopy (deblurring, denoising, and optical deconvolution) provides researchers with detailed information on samples. Recent general-purpose deep learning solutions advanced in this task. Yet, without consideration of the underlying physics, they may yield unrealistic and non-existent details and distortions during image restoration, requiring domain expertise to discern true features from artifacts. Furthermore, the large expressive capacity of general-purpose deep learning models requires more resources to train and use in applications. We introduce DeBCR, a physics-informed deep learning model based on wavelet theory to enhance microscopy images. DeBCR is a light model with a fast runtime and without hallucinations. We evaluated the image restoration performance of DeBCR and 12 current state-of-the-art models over 6 datasets spanning crucial modalities in advanced light microscopy and cryo-electron tomography. Leveraging optic models, DeBCR demonstrates superior performance in denoising, optical deconvolution, and deblurring tasks across both LM and cryo-ET modalities.

## Introduction

Microscopy is essential in biological research. In light microscopy (LM), fluorescence microscopes^1,2^ are fundamental for examining cellular and tissue structures. In electron microscopy (EM), cryo-electron tomography^3^ (cryo-ET) uniquely enables the analysis of natively preserved biological samples *in situ* at the Ångström level. However, limited photon/electron doses during imaging often result in noisy images with artifacts and limited resolution. In LM, the limited photon budget necessitates balancing spatial resolution, temporal resolution, and imaging duration. In cryo-ET, similar considerations apply to the electron budget in the tilt series: electrons are required to form a contrast, however, excessive dose destroys the sample^3^. A trade-off between the image quality and the acquisition costs (e.g. sample damage, longer imaging time, and more complicated data processing, etc.) occurs in both the LM and cryo-EM/ET imaging. Consequently, achieving higher-quality, interpretable images has been a longstanding focus in both LM^4^ and cryo-EM^5,6^. In super-resolution (SR) fluorescence microscopy, resolving point sources in imaging devices can be improved either by redesigning hardware configurations or through image processing. For LM, light-sheet microscopy^7^, structured illumination microscopy^8^ (SIM), and stochastic optical reconstruction microscopy^9^ (STORM) offer significantly advanced image resolution. In cryo-EM^10–12^, single-particle analysis^13^ (SPA) and subtomogram averaging (StA)^3^ allow for a resolution improvement via averaging over multiple copies of the same molecular type. However, these resolution enhancement solutions often require expensive advanced hardware and skilled personnel. Balancing the cost of technological advancements with image quality across various imaging modalities is important for their rapid and effective application.

Image processing^14^ and deep learning^15^ (DL) demonstrate exceptional performance in computer vision. Under the general topic of ‘image enhancement’, various DL approaches have succeeded in restoring high-resolution details from noisy inputs. DnCNN^16^ utilizes a residual structure to effectively remove Gaussian noise. RCAN^17^ combines deep residual structures with the channel attention mechanism for superior results. MPRNet^18^ employs multi-stage learning strategies and brings deblurring/denoising to new levels. For generative models, ESRGAN^19^ enhances image quality by adapting the generative adversarial model for image enhancement. DDPM^20,21^ improves image quality by generating new signals from known distributions (often Gaussian distribution). These models require noise-free or high-resolution images as ground truth (GT). When GT images are unavailable, the Noise2Noise^22^ (N2N) model offers a solution by denoising images without the corresponding GT.

Specific to biology, many studies^23,24^ have embraced DL to enhance performances in image-based applications. In LM, DL excels in segmentation^25^, deblurring^17^, and denoising^26^. For cryo-ET data, DL models are used for denoising^27^, compensating the “missing wedge”^28^, and accounting for the conformational flexibility of proteins^29^. Specifically, researchers have developed DL toolsets tailored for microscopy. The U-Net^30^ model, with its downsampling/upsampling structure, is versatile and performant in medical image segmentation. Furthermore, derivative models with improved structure and regularization, show promise for widefield microscopy image restoration^25^. CARE^31^ model excels in denoising, super-resolution, and 3D isotropic restoration. UniFMIR^32^ proposes large pre-trained models and fine-tunes them for various microscopy image tasks. In cryo-ET, traditional denoising solutions^33^ like Gaussian and median filters preserve information only within specific frequency ranges. High-frequency signals in cryo-ET images are broadened and convoluted^34,35^ with noise, making blind noise removal challenging. DL-based solutions like Topaz^36^ and cryo-CARE^27^ effectively remove noise while preserving signals. Biological research increasingly relies on deep neural network (DNN) models for denoising, deblurring, and optical deconvolution tasks. However, researchers often increase the model size to achieve better results. From CARE to ESRGAN, the number of trainable parameters has increased 150 times, demanding more computational resources and longer training times, and posing challenges in regularizing outputs. Larger models can generate unrealistic patterns, resulting in hallucinations in microscopy restorations.

Here, we propose a physics-informed DL model, DeBCR, for denoising, deblurring, and deconvolution tasks in microscopy images. The physics process for imaging can be represented as 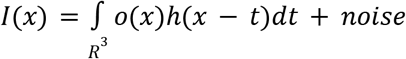, where the sample information *o*(*x*) is convolved with the kernel *h*(*x* − *t*)^37–39^ to produce the image *I*(*x*). Using the Beylkin-Coifman-Rokhlin^40,41^ (BCR) wavelet theory, we implemented this process within a DNN structure^42^ as DeBCR. For details, see the ‘Microscopy imaging model’ section in ‘Online Methods’. To evaluate our model, we compared DeBCR to 12 state-of-the-art (SOTA) models (from DnCNN to UniFMIR for LM; Topaz and cryo-CARE for cryo-EM) across six real microscopy datasets. These experiments assess denoising, deblurring, and deconvolution tasks. DeBCR consistently ranked highest in all assessments. It achieved the best metrics and lowest hallucinations in LM experiments and showed clear advantages in cryo-ET experiments. Leveraging physics backbones, DeBCR uses ∼210 times fewer trainable parameters (ESRGAN) and runs ∼480 times faster (ESRGAN) than others. This work highlights the benefits of using physics models to guide the design of DNNs and provides a new tool for image restoration.

## Results

In LM, obtaining high signal-to-noise ratio (SNR) images remains a general challenge. Fluorescent microscopy entails diverse requirements during imaging: spatial/temporal resolution, phototoxicity, imaging depth, etc. to produce high SNR images. Higher SNR can be achieved by increasing the illumination power or extending the exposure time of the samples. Yet, maximizing all these factors simultaneously is often unfeasible due to photodamage. In cryo-ET^10–12^, the contrast of tomograms is capped by the limited electron dose^3^, however, a higher electron dose leads to sample damage^43^. A trade-off between the image quality and the acquisition costs (sample damage, longer imaging time, and more complicated data processing, etc.) occurs in both the LM and cryo-EM/ET imaging.

Here we address signal restoration in LM and cryo-ET using DeBCR (see Methods) using four benchmarking LM datasets and two cryo-ET datasets. Fig.1a and Fig.S1a,b presents the neural network structure of DeBCR. Based on the previously proposed BCR approach^44^, we approximated the image-enhancement operator with two components: integral operators (blue panel) and pseudo-differential operators (green part). The integral operators comprise subsets of 2D convolutional layers, while the pseudo-differential operator corresponds to the BCR-inspired residual convolution layers. For further details, please refer to the method section in the supplementary material, as well as Fig.S1c and Fig.S2.

**Fig. 1.**
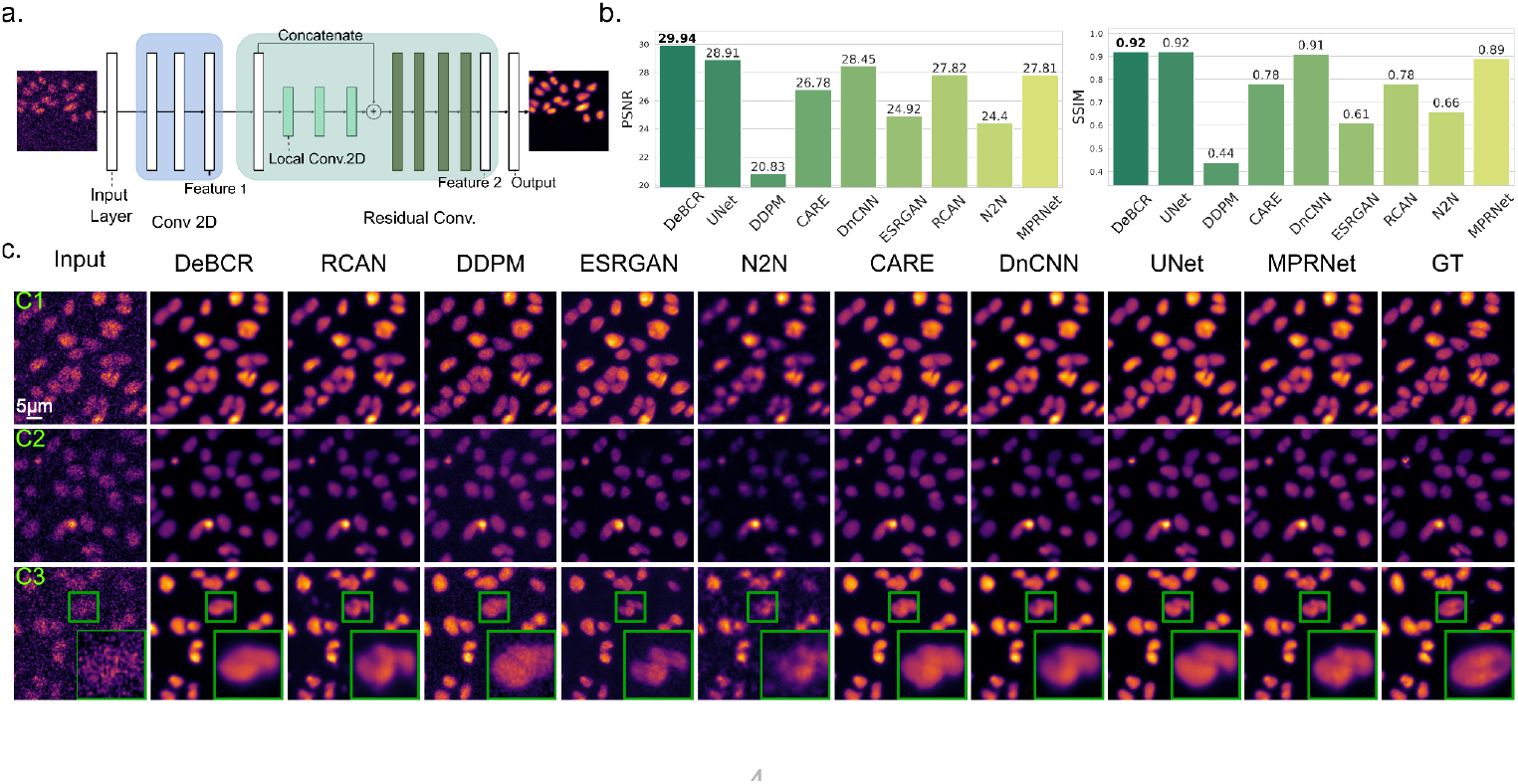
Denoising 2D confocal microscopy images of the flatworm *Schmidtea mediterranea*. The dataset includes images of noisy input alongside their corresponding ground truth (GT) obtained by longer exposure time and higher laser intensities. The noise levels are categorized according to the 3 illumination levels (C1 - medium, C2 - weak, C3 - extremely weak). **a**, The structure of the DeBCR models. This model consists of two parts. The blue portion corresponds to the forward operator (refer to the supplementary material), primarily involving conv2D operations. The green part pertains to the pseudo-differential operators, mainly comprising the BCR-inspired residual deep neural network (DNN) structure. **b**, Evaluation by SSIM↑/PSNR↑ on the test images. The highest ranking is marked with bold font. **c**, Comparison of the denoising performance to the to 8 SOTA denoising models. ROI in C3 shows zoom-in for detailed restorations.

### Denoising of 2D confocal microscopy images

Originating from CARE^26^, this dataset contains confocal 2D fluorescence microscopy images of the flatworms *Schmidtea mediterranea*. Due to the extreme sensitivity of the worms to the illumination levels, even normal laser intensity can induce flinching of the samples. The dataset contains images from 3 laser power levels (C1 - medium, C2 - weak, C3 - extremely weak) and the GT. As the laser power decreases, noise becomes increasingly dominant. We compared DeBCR for this 2D denoising task to 8 SOTA models (CARE^26^, RCAN^17^, DnCNN^16^, DDPM^20^, ESRGAN^19^, N2N^22^, U-Net^25^, and MPRNet^45^). To be noticed, the U-Net here referred to the advanced U-Net structure designed for brightfield microscopy segmentation^25^, rather than the originally proposed^30^. To quantify performance, we computed the metrics Structural Similarity Index Measure^46^ (SSIM) and Peak Signal-to-Noise Ratio^46^ (PSNR) for the models. DeBCR achieved the highest performance (Fig. 1b) with 29.94 dB/0.92 (PSNR↑/SSIM↑). Following DeBCR, the advanced U-Net (28.91 dB/0.92) and DnCNN (28.45 dB/0.91) rank second with a noticeable gap to DeBCR. Despite its high performance in SNR, DDPM ranked lowest due to the salt-pepper-like artifacts. All models demonstrated satisfactory performance (Fig. 1c) by restoring signals from the noisy inputs for C1 and C2 intensities. The restoration performance varies as the signal reduces to the extremely weak level of C3. Here, the N2N failed to restore the cells’ morphology. Even though other candidates restored high-resolution signals, they all exhibited distortions and generated non-realistic patterns. In the region of interest (ROI), ESRGAN misconstructed the input pattern as two cells, while there is only one cell in GT. DDPM’s restoration exhibits strong salt-pepper artifacts, while DeBCR uniquely removed the noise without introducing artifacts and maintained a high similarity to the GT. Furthermore, compared to a large model like UniFMIR DeBCR generates far fewer hallucinations (see spplement and Fig. S3).

### Denoising of 3D confocal microscopy data

Confocal microscopy enables optical sectioning without physically slicing the sample^47^. This allows the 3D imaging along the axial direction and potentially live. Yet, optimizing SNR, acquisition time, and specimen viability in 3D confocal microscopy requires extending the photon budget^26^. Furthermore, the confocal images have worse axial resolution due to the light diffraction. Therefore, restoring axial details poses even greater challenges in this task. Introduced by CARE, the *Tribolium castaneum* embryos dataset contains the confocal microscopy image stacks in low- and high SNR allowing us to extend the photon budget computationally. Similarly to the CARE, we adopted the high SNR data as a GT and trained all models for 3D denoising. Fig. 2a depicts the reconstructions of DeBCR in both lateral and axial planes. With the inputs in extremely low SNR, DeBCR effectively restores signals with both high axial- and lateral resolutions. A detailed inspection of ROIs in Fig. 2b for the 3D restorations demonstrates the performance of the models. N2N, designed primarily for denoising, exhibits a limited ability to restore corrupted signals. The generative models DDPM and ESRGAN introduce texture-like artifacts, as reflected in inferior metrics assessments PSNR↑/SSIM↑ (Fig. 2c). DnCNN, U-Net, CARE, and RCAN wrongly restored the non-existent patterns, while MPRNet lost some of the patterns in GT. DDPM restored the signal correctly but with poor metrics. DeBCR stands out uniquely for restoring high-resolution signals and keeping them identical to the GT without artifacts. In Fig. 2c, DeBCR achieved the best performance among all candidates with PSNR↑/SSIM↑ as 20.24 dB/0.62. This demonstrates the reliability of DeBCR in denoising anisotropic 3D LM data. As a reliable restoration toolset, DeBCR enables the confocal imaging process to be conducted faster, with higher time resolution and lower photodamage, which is advantageous for live imaging.

**Fig. 2.**
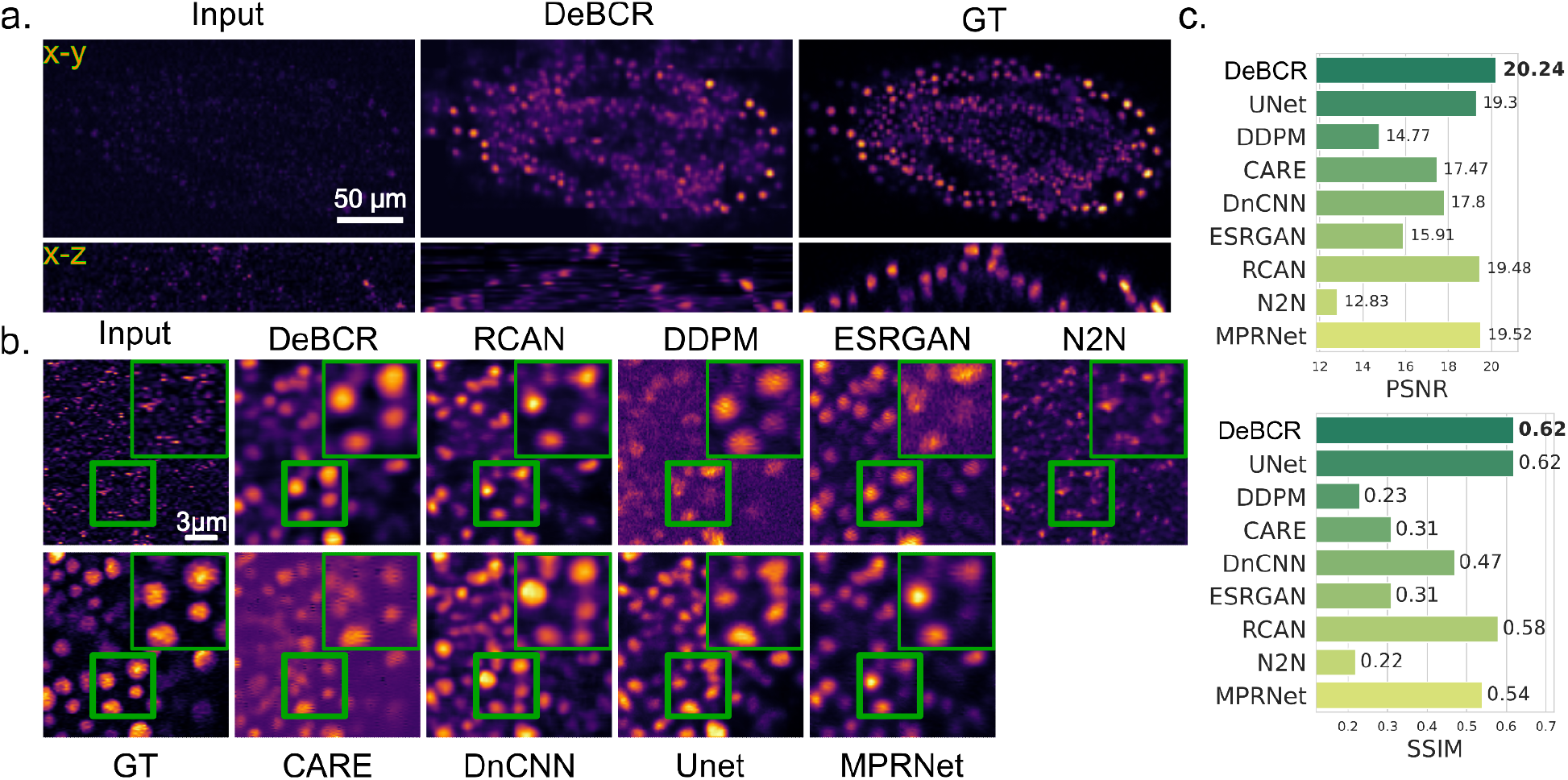
3D Denoising on the confocal microscopy datasets of *Tribolium castaneum* embryos. **a** The input images exhibit low SNR. DeBCR’s reconstruction in xy- and xz- planes closely matches the GT. The xy plane restoration captures most details similar to GT. **b**, The comparison of ROI to other candidates’ reconstructions. The low SNR inputs elevate the task from mere denoising to denoising and compensating for corrupted signals. DeBCR uniquely restored the accurate patterns in GT without introducing new artifacts. **c**, Metrics evaluation in PSNR↑/SSIM↑ for all restorations.

### Recovering SIM-like signal from widefield LM data

In advanced SR microscopy, achieving high spatial and temporal resolutions while maintaining specimen viability hinges on managing the photon budget effectively. Data-driven SR microscopy methods allow researchers to extract more information from specimens despite low SNR. By leveraging paired training data, these methods can enhance low-resolution images to high-resolution using DNN models. This approach is particularly appealing as low-resolution images can be obtained quickly and affordably through methods like widefield microscopy. However, a frequently overlooked issue is the occurrence of hallucinations during DNN-based reconstruction. Many SOTA models lack domain-specific constraints, leading them to generate fake patterns (referred to as hallucinations in this context) that do not correspond to GT data. This problem is particularly critical in research microscopy, where identifying hallucinations can be challenging even for domain experts. To validate DeBCR’s performances in SR, we utilized a microscopy dataset featuring *S. aureus* from the TAGAN^48^ baseline model. This dataset includes image pairs captured using widefield microscopy and Structured Illumination Microscopy^47^ (SIM). SIM images serve as GT for assessing restorations from blurry widefield images. After training, we compared all the restoration results against DeBCR (Fig. 3a). The N2N model, designed primarily for denoising tasks, performs sub-optimally. While other candidates successfully restore SR details, they exhibit varying levels of artifacts and distortions. Upon closer inspection of the ROI, RACN, TAGAN, DDPM, ESRGAN, and U-Net introduce fake patterns during resolution enhancement. MPRNet effectively controls hallucinations but shows inferior resolution restoration. Only DnCNN and DeBCR demonstrate comparable performance in artifact compression and high-resolution detail restoration. Fig. 3b presents the evaluation metrics in terms of PSNR↑/RMSE↓. SSIM^49^ evaluation requires well-aligned images, which fails in this case due to sample drift between the microscopy images. For the assessments, DeBCR achieves top rankings with 29.94 dB/0.04. U-Net and DnCNN perform similarly, securing the second position. In contrast, N2N ranks lowest in the list.

**Fig. 3.**
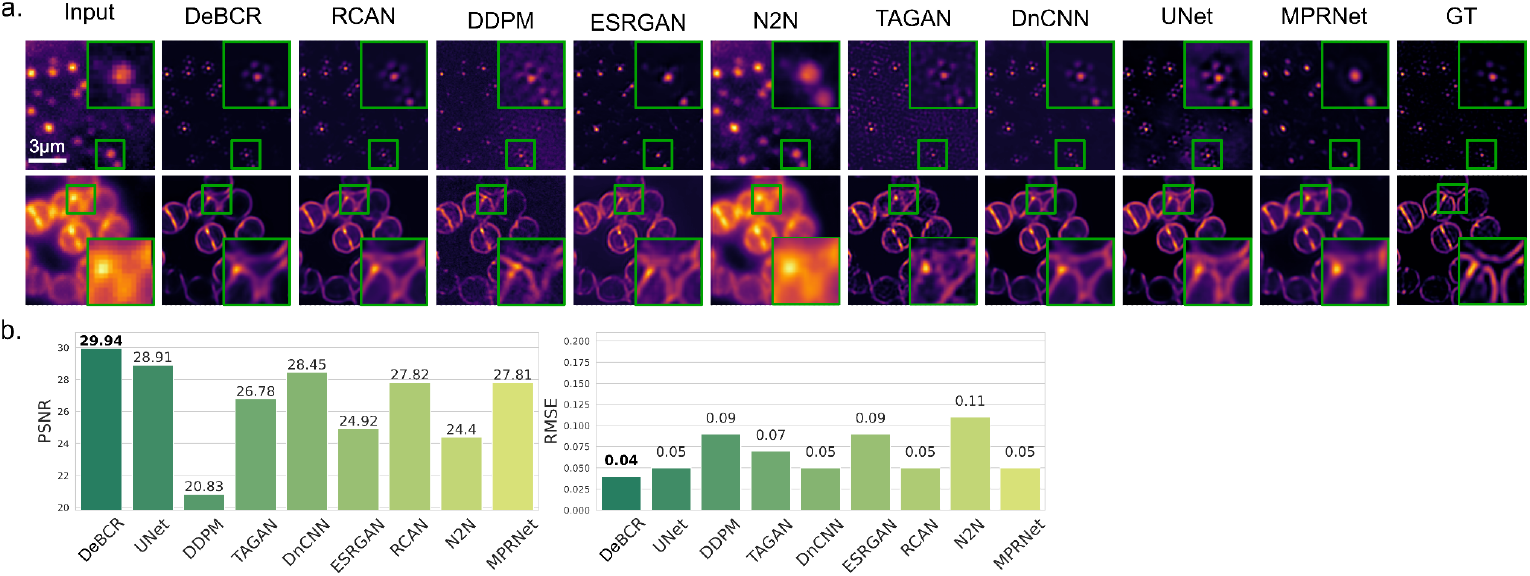
Optic deconvolution on brightfield microscopy images for *S. aureus*. The dataset comprises pairs of images captured using brightfield microscopy and Structure Illumination Microscopy (SIM). The SIM images serve as pseudo-ground truth (GT) for optic deconvolution. We compared our results to TAGAN as a baseline. **a**, visualization of selected restoration results. **b**, the evaluation metrics PSNR↑/RMSE↓ for the test results.

### Recovering STED-like signal from confocal LM data

STimulated Emission Depletion Microscopy^47^ (STED) requires advanced hardware to provide high-resolution images. This SR dataset of *F-Actin* contains both confocal microscopy images and their well-registered STED images. We tested DeBCR to recover an STED-like signal using confocal microscopy. This SR experiment was configured by TAGAN^48^, we adopted TAGAN as the baseline as well. In Fig. 4a all models, except N2N, achieved overall good performance during the SR task. The tested generative models ESRGAN, DDPM, and TAGAN hallucinated by restoring fake patterns that do not exist in the GT. Compared to others, DeBCR effectively enhances resolution without hallucinations. We quantitatively assessed the model performance using PSNR↑/RMSE↓. DeBCR achieved the best performance at 27.01 dB/0.05 (Fig. 4b). While TAGAN also showed a similar performance to DeBCR, its restoration is characterized by a high percentage of hallucinations.

**Fig. 4.**
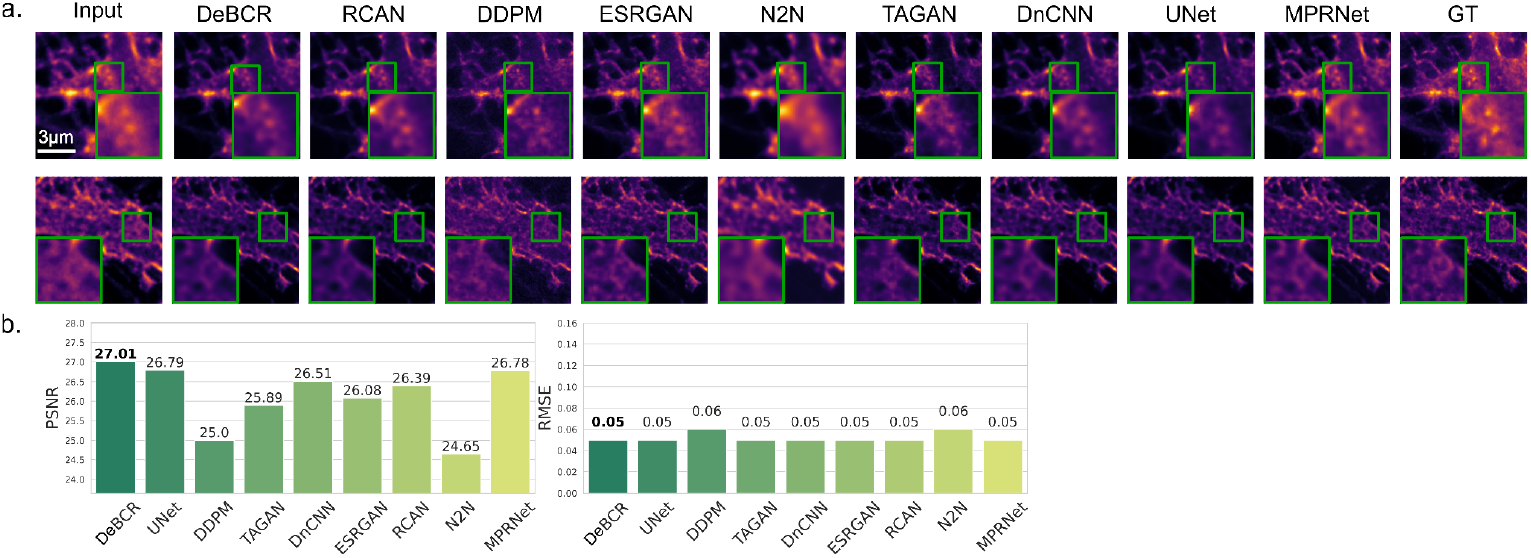
Recovering STED-like signal from confocal microscopy images for *F Actin*. This dataset contains pair images from confocal and STED^43^ images for *F-actin* samples. The STED serves as the pseudo-GT. **a**, visualization of the results after the restoration. **b**, quantitative the evaluation with PSNR↑/RMSE↓. DeBCR achieved the best performance with PSNR/RMSE scores of 27.01/0.05, ranks at the top position in the comparison.

### Restoring the low-frequency signal by cryo-ET data

In cryo-EM, the signal is generated by defocused phase contrast. This results in low contrast transfer for low frequencies^50^. Due to the samples’ low radiation tolerance, biological applications limit the electron dose for cryo-ET to 100-150 e-/Å^2^ distributed over 40-60 tilted images. This weak electron dose leads to dominant noise^51^. Additionally, the limited angular range of ±60 degrees during tilt series collection causes lower anisotropic resolution in the direction of the electron beam, leading to the “missing wedge” of information in Fourier Space. These constraints result in cryo-ET volumes with low SNR.

In this experiment, we performed denoising and signal recovery on the publicly available Tomo110 dataset from the cryo-CARE report^27^, which contains raw cryo-ET data of *Chlamydomonas reinhardtii* cilia. For cryo-ET denoising applications, where ground truth (GT) data is unavailable, we applied the N2N^22^ framework for DeBCR (Fig. 5a). Specifically, we split the noisy movies of tilt series into two groups (odd and even frames), performed two statistically independent tomographic reconstructions and trained the DeBCR model to transfer one half to another. We compared our denoising results with commonly used methods in cryo-ET: Cryo-CARE, Topaz^36^, and a Gaussian^36^ filter (Fig. 5b,c). Visual analysis showed that DeBCR outperformed the other methods in denoising microtubules and membranes. While Gaussian filters primarily affect low-frequency signals and tend to lose high-frequency information, DL-based methods like Topaz, Cryo-CARE, and DeBCR demonstrated superior noise reduction with high-frequency details. To quantify performances, we calculated the Fourier Ring Correlation (FRC) between the denoised even- and odd volumes of the test dataset (Fig. 5d). FRC evaluation indicated that DeBCR restored a significantly larger amount of information across all frequencies compared to the other methods. Furthermore, by analyzing even/odd volumes, we estimated the noise spectrum^52^ from the difference between two groups of volumes (see Supplementary materials, Fig. S4). This approach revealed a clear noise reduction effect of 20-40% when using the DeBCR model compared to raw data (Fig. 5e, raw vs. DeBCR). Overall, these results highlight the effectiveness of the DeBCR model in cryo-ET denoising tasks.

**Fig. 5.**
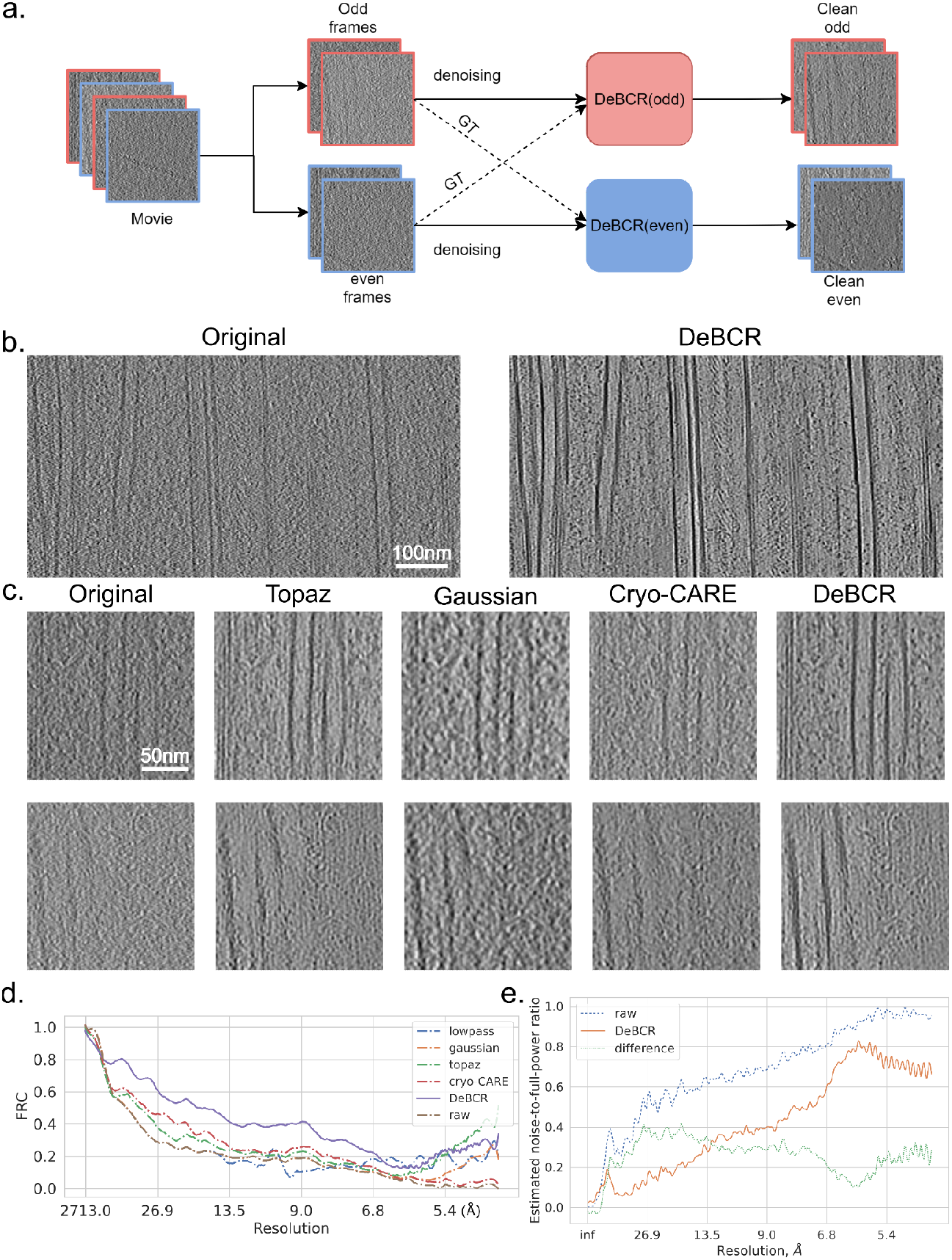
Denoising cryo-ET data. **a**, In the absence of ground truth data, we integrated DeBCR into the Noise2Noise^22^ framework. The model is trained to learn the transfer mapping between odd and even frames within one tomograph. This enables the model to denoise the dataset without GT. **b**, Denoising the tomogram containing microtubules and membranes by DeBCR. DeBCR preserves the contrast of the target features while effectively filtering out the noise. **c**, Comparison of the denoising methods for DeBCR and SOTA denoising models in cryo-ET. **d**, Fourier Ring Correlation (FRC) evaluation for the restorations. DeBCR ranks at the top in all frequency ranges with higher FRC values. **e**, Estimated spectral noise-to-full-power ratio before and after the DeBCR-based restoration.

### Restoring the high-frequency signal by Cryo-ET data

We next challenged DeBCR to the dataset containing a membrane protein RyR1 is imaged in native membrane vesicles extracted from rabbit muscle^53^ (EMPIAR-10452). Here, subtomogram averaging of a RyR1 was performed and we could use it as ground truth to evaluate the high-frequency signal restoration. We adopted the same N2N framework combined with DeBCR for this experiment. Fig. 6a shows the restoration results in X-Y and Y-Z planes. DeBCR effectively suppresses noise in both dimensions. Interestingly, by analyzing the X-Z views we realized that in DeBCR-recovered tomograms it is possible to identify the sample-vacuum interfaces, which are hard to distinguish in the unfiltered tomograms. Fig. 6b provides a detailed examination of ROIs of restorations. In contrast to the noisy inputs, the denoised outputs highlight the membrane protein RyR1 along the membrane by effectively removing noise. The FRC evaluation in Fig. 6c indicates the signal enhancement across nearly the entire frequency range, indicating higher internal consistency.

**Fig. 6.**
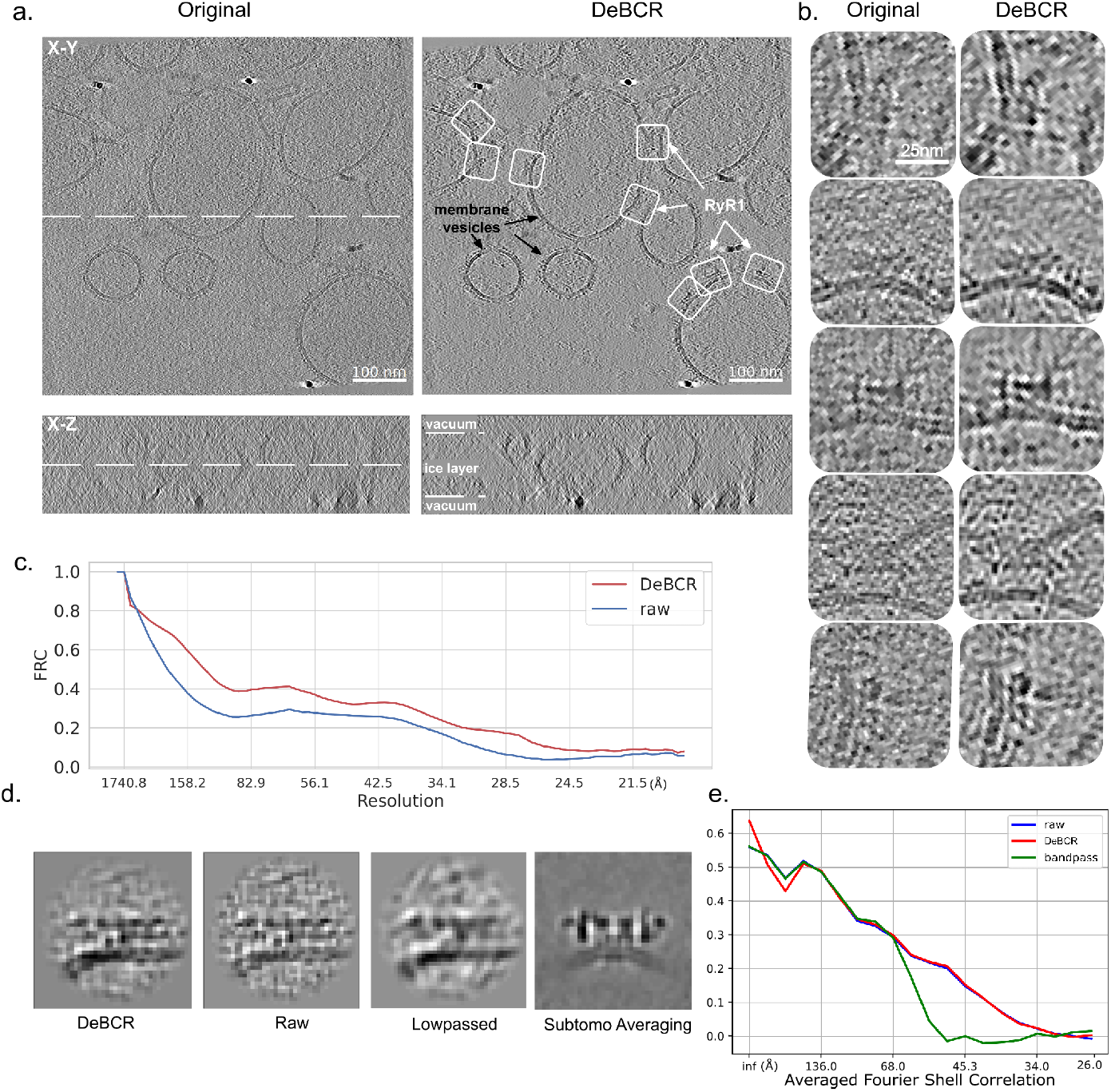
Denoising cryo-ET tomograms of fractions of rabbit skeletal muscle containing a membrane protein RyR1. Due to the missing GT, we employed the Noise2Noise^22^ framework for the denoising task for denoising. **a**, Denoising for the tomogram in the central X-Y and X-Z planes (white dashed lines denote the corresponding orthogonal views). DeBCR successfully eliminates noise from the original tomogram while preserving high-frequency information. The X-Z plane illustrates the restoration in the axial direction. The examples of membrane vesicles (black arrows) and membrane protein RyR1 (white boxes) are annotated. **b**, Comparison of selected patches of the membrane protein RyR1 from the original and the DeBCR-denoised tomogram shown in **a**, from the X-Y planes. **c**, FRC evaluation of the DeBCR denoising results. In comparison to the inputs, DeBCR effectively restores a greater amount of information across all frequency ranges. **d**, Individual subtomograms containing RyR1 from one to three from the left show the same particle cropped from DeBCR denoised, raw, and low-passed tomograms. The fourth panel shows the subtomogram average of RyR1 used as the ground truth for calculating FSC in panel e. **e**, Averaged Fourier Shell Correlation of raw, bandpassed, and DeBCR-denoised individual particles to the ground truth.

We probed if the recovered signal contained high-resolution information. We identified RyR1 particles and their orientations to the average in the tomograms of the test set using template matching, manually inspected them, and excluded false positives. We calculated 3D Fourier Shell Correlation (FSC) between the subtomograms containing the receptors and the subtomogram average (Fig.6d,e). The average FSC for raw and DeBCR denoised data is similar, while a band-pass filter (equivalent to a Gaussian filter) reduced this FSC. This measurement demonstrates that DeBCR suppresses the noise without deterioration of the signal. Denoised outputs of DeBCR, can be used to generate preliminary reconstructions, as shown in Fig.S5.

## Discussion

In this study, we introduce DeBCR, a physics-informed DNN toolkit tailored to the denoising, deblurring, and deconvolution of microscopy images. Our model DeBCR draws inspiration from the optical convolution process. Consequently, it has the advantage of effectively constraining artifacts during the reconstruction process while utilizing fewer trainable parameters. We show that this makes DeBCR highly robust and results in enhanced performance of deconvolution, debluring, and denoising while avoiding hallucinations. Uniquely, DeBCR can efficiently process diverse data ranging from various fluorescent microscopy modalities to cryo-ET data.

We demonstrated in several computational experiments that DeBCR recovers the high-resolution or high-SNR signal and avoids hallucinations and fake patterns during the restoration. In the LM experiments, our model consistently outperforms other SOTA models (from DnCNN, MPRNET to CARE, etc.) in terms of metrics PSNR↑, SSIM↑, and RMSE↓. In the cryo-ET experiments, DeBCR emerges as the top performer among other candidates (Topaz, cryoCARE, and bandpass denoisers). It successfully restored signals across the entire frequency spectrum, as we show by the Fourier Ring Correlation (FRC) evaluations. In the global view, our model outperforms 12 SOTA models for the tasks of deconvolution, denoising, and deblurring. This offers Life Science researchers a unique generalist go-to model and allows them to extract better information from microscopy images.

The efficiency of DeBCR has only been extensively demonstrated in the context of restoration tasks based on theoretical foundations such as denoising, deblurring, and deconvolution. Its usage in other applications needs to be further investigated and adapted. The limitations of DeBCR to date are in its applicability to more specific microscopy image tasks including surface projection^26^, isotropic reconstruction^32^, and direct segmentation^48^. Furthermore, being a relatively small model, retraining of DeBCR on user-specific data is required. For tasks such as projection and isotropic reconstructions, DeBCR may not be an optimal solution due to constraints stemming from both theoretical limitations (a governing physical model must be known) and the number of trainable parameters. Further adaptation of the DeBCR structure to encompass a broader spectrum of microscopy image restoration tasks is anticipated in future research.

In summary, our experiments demonstrated that integrating physics-based models to inform the design of DNN models can significantly enhance efficiency and reliability in the microscopy image restoration process without the introduction of non-existent patterns. The DeBCR model holds promise for numerous downstream applications in microscopy, enabling lower laser intensity, shorter imaging times for the same contrast and resolution, or higher contrast without signal degradation for cryo-ET data.

## Data and Code Availability

The TensorFlow model is provided in the open-source GitHub repo (https://github.com/leeroyhannover/DeBCR). For quick deployment, we offer access to our model and weights in the CodeOcean capsule (https://codeocean.com/capsule/9904237/tree/v1) to reproduce the test results in this paper. The project in CodeOcean covers all the experiments of our DeBCR model in this paper including the test dataset and trained weights. The full version of 6 datasets (training+validation+testing) is released on Zenodo (https://zenodo.org/records/12626122). To reproduce the training process of DeBCR, we offered a demo tutorial to prepare the training process as a Google Colab notebook (https://githubtocolab.com/leeroyhannover/DeBCR/blob/main/notebooks/DeBCR_train.ipynb).

To benefit the readers for their own project using our model, we will release a comprehensive tutorial from data preparation to training/testing in Colab.

## Acknowledgements

This work was partially funded by the Center for Advanced Systems Understanding (CASUS) which is financed by Germany’s Federal Ministry of Education and Research (BMBF) and by the Saxon Ministry for Science, Culture, and Tourism (SMWK) with tax funds on the basis of the budget approved by the Saxon State Parliament. MK was supported by the Heisenberg award from the DFG (KU 3222/2-1), as well as funding from the Helmholtz Association. The authors thank HelmholtzAI (grant tomoCAT).

## Author contributions statement

RL, AYu, XC, MK and AY conceived the idea and planned the computational experiments. RL designed the mathematical models, RL, AYu, XC wrote the program code and conducted computational experiments. AYu, XC prepared and acquired cryo-ET experimental data. RL, AYu, XC, MK and AY wrote the manuscript.

## Online Methods

### Microscopy imaging model

The imaging process in fluorescence microscopy and cryo-EM can be described by the simplified microscopy imaging model below. The sample information *o*(*x*) goes through the optical lens system *A*(*t*)^38^ and contributes to the contrast of the final images *I*(*x*). The *A*(*t*) is defined as the convolution operator based on the different physics configurations (e.g. hardware, illumination, processing, etc.). In fluorescence microscopy, *A*(*t*) corresponds to the Point Spread Function (PSF), whereas it corresponds to the Contrast Transfer Function (CTF) - the PSF in Fourier space - in cryo-EM. Due to variations in physical models, the *A*(*t*) assume diverse forms across various microscopies. In widefield microscopy, it can be simplified as 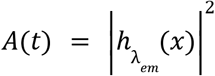, taking into account emission amplitude distribution 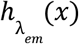, while in confocal microscopy it takes a distinct form as 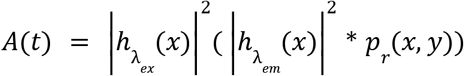, additionally considering excitation amplitude 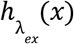.The function *p*_*r*_(*x, y*) represents the pupil function specific to confocal microscopy, reflecting hardware differences attributed to the presence of a pinhole. An image *I*(*x*) represents a measurement of genuine information *o*(*x*) through the convolution process *I*(*x*) = *A*(*t*) * *o*(*x*). In reality, the limitations of the optic models and the hardware precision introduce noise/artifacts ε into *I*(*x*) as in equation (1). Both the convolution and noise terms ε account for the blur and noise effects in final microscopy images. Restoring of the *o*(*x*) from the acquired *I*(*x*) holds the key to reveal the high-resolution samples information. General computer vision tasks encompass image-quality enhancement tasks (deconvolution/denoising/deblurring) without specific consideration for their underlying physics. From the physics perspective, the pursuit of the inverse operator *A*(*t*) ^−1^ stands for optic deconvolution^54^. Removing the disturbing item ε refers to the task of denoising. Deblurring generally includes both deconvolution and denoising, as the blur effect in images can result from both noise (e.g., Gaussian noise) and convolution.

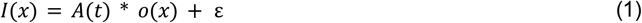

Depending on the microscopy type, the convolution operator *A*(*t*) ranges from the point spread function^37,38^ (PSF) in fluorescence microscopy to the Contrast Transfer Function^39^ (CTF) in cryo-EM. Denoising tasks refers to removing the item ε. Deconvolution tasks indicate the inverse of the operator *A*(*t*) to restore the *o*(*x*). Deblurring stands in general the combination of both previous tasks. Many of the traditional deconvolution solutions^2,54^ concentrate on searching for the inverse form *A*(*t*)^−1^. Yet, the process *A*(*t*)^−1^ is ill-posed^55^ due to the contribution of ε. This instability in the solution often leads to artifacts and distortions. Besides, this inverse process requires the posterior of PSF/CTF for every image. The PSF/CTF correlates to various physical parameters (e,g. wavelength, Airy disc size, and diffraction ratio, etc.). Determining these parameters in most cases can be challenging. Thus, the PSF/CTF information is generally missing. The deconvolution solution without PSF/CTF is defined as blind deconvolution^56^. DL-based blind deconvolution^21,57^ solutions have demonstrated their potential. However, those methods still need to simulate the artificial PSF/CTF used during imaging and then conduct the inverse of convolution. In this work, we introduce a physics-informed toolset DeBCR tailored for microscopy image restorations. Based on the optical model, this model is specifically designed to address denoising, deblurring, and deconvolution tasks without explicit CTF/PSF parameters.

We denoted the previous imaging process further as the compact of the integral operators 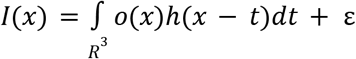. The Beylkin-Coifman-Rokhlin (BCR) wavelet theory^40,41^ proved that the integral operator can be approximated with a combination of wavelets at low computational costs. This concept inspired the work for transferring the wavelet approximation of the integral operator to the basic DNN structures^42^. Applying this to the imaging process in the stepstone work^44^ we demonstrated the strength of the physics-informed deep neural network. This BCR-based model provided an innovative approach to address the task of blind deconvolution. Here, we adopted the m-rBCR model as a baseline and proposed the general method of DeBCR. We extended the scenarios of m-rBCR from deconvolution to denoising and deblurring tasks in both LM and cryo-EM applications. This work highlights the benefits of incorporating physics models to guide the design of deep neural networks (DNNs).

## DNN representations

The process of obtaining the *o*(*x*) from the measurements *I*(*x*) can be denoted as pursuing the inverse mapping *I*(*x*) → *o*(*x*). A general format of inverse problems^58^ can be denoted as in equation (2) below. The *b* is the edge measurement from the true signal *u*^*^, the *A* stands for measurement operators - the optical convolution in this work. The process in equation (2) is referred to as the forward process, which forms the microscopy images *b* from the sample information *u*^*^.

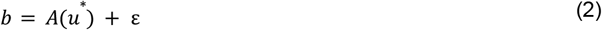

As discussed in the seminal work^44^, the solution to the inverse problem^58^ is available by optimizing the item *min*|*A*(*u*) − *b*|^2^. Yet, the *b* in microscopy contains the dominant noise term ε in the real world. This hinders the unique solution *u*^*^ through the traditional methods. The mapping is neither unique nor stable. Such a problem is defined as the ill-posed inverse problem^55^. Due to the ill-posed nature of the problem, optimization pursuits lack robustness. To stabilize the optimization solution, a general trick is to add the regularization term ϕ(*u*)^59^ as in equation (3). This regularization term mitigates non-uniqueness challenges. The traditional regularization strategies contain the sparsity regularizer^60^ and variation regularizer^61^.

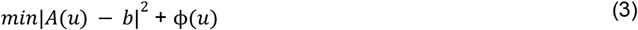

In contrast to classical regularization strategies, the cornerstone work^44^ showcased the regularization method in DNN’s structural design by employing the Beylkin, Coifman, and Rokhlin (BCR)^41^ decomposition for microscopy imaging models. The inverse problem can be decomposed into the operation of two components in equation (4). This approximation involves the combination of two operators: the integral operator *K*^*T*^ and the pseudo-differential operator *B* = (*K*^*T*^ *K* + ε*I*)^−1^.

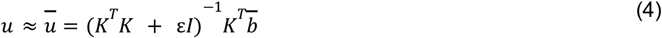

Introducing the equation (4) to equation (1) induced the form of the inverse operator *A*^−1^ as the equation (5) below. The computation of equation (5) yields the solution 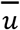, which is the approximation of the genuine information *u*^*^.

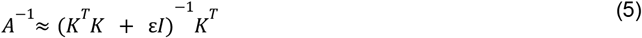

Validated in the prior work^44^, the integral operator can be represented with wavelet approximation based on BCR decomposition. This approximation has a natural connection to the matrix operation in DNN. For more details on the BCR neural networks, please refer to Fig.S1 in supplementary materials.

## Experiments configurations

The dataset in all experiments is split into training/validation/testing phases with the ratio 0.8/0.1/0.1. Benefiting from the backbone of BCR theory and microscopy imaging models, DeBCR operated effectively without the specialized optimizer, learning rate, or training strategies. We used Adam optimizer with a learning rate of 0.001 from TensorFlow. We settled the training epochs as 10^5. Due to the use of early-stopping to avoid overfitting, the training process converged generally before the 10^5 depending on the task types. Since the model contains the backbone of microscopy imaging theories, it does not need the extra regularization item for loss function. The loss function we utilized for DeBCR is Mean Square Error (MSE) on the reconstruction and ground truth (GT) images. The specific calculation equation is listed below. All experiments are conducted on the Tesla V100 GPU.

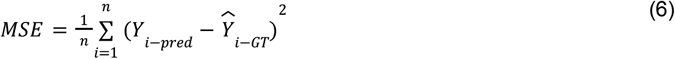

### Used metrics

To evaluate the performance of the learning process, we utilized four metrics^62^ for the experiments. Specifically, PSNR, SSIM, and NRMSE were employed for the experiments in light microscopy, while the 3D Fourier Shell Correlation^63^ (FSC) and the 2D version Fourier Ring Correlation^64^ (FRC) were utilized for the cryo-electron tomography (cryo-ET) experiments. We abstained from using PSNR, SSIM, and NRMSE for the cryo-ET tasks because these experiments lack ground truth data, which is required by these metrics for calculations. The specific equations for the metrics are listed below.

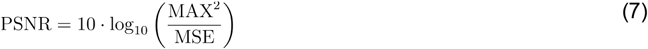

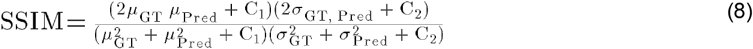

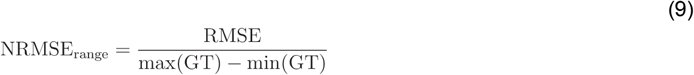

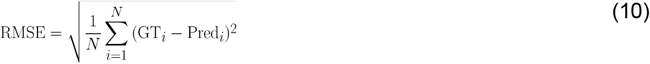

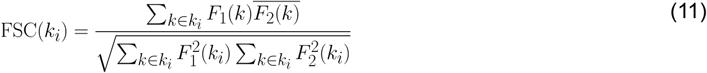

In equation (10) where k_i_ is the single-frequency 2D ring (in FRC) or 3D shell (in FSC); F_1_ and F_2_ are radially averaged Fourier transforms of the two correlated images (in FRC) or volumes (in FSC), containing the same signal, but independent noise. The even/odd datasets and independent particle half-set averages served us as such same-signal noise-independent data pairs.

### Parameters and runtime comparison

We evaluated the number of trainable parameters (in millions) and the running time (in seconds) in testing to highlight the advantages of employing slim models with physical backbones. The results are presented below in Table 1. The time efficiency aligns with the principle that larger models consume more time for calculations, limiting data and application throughput. DeBCR achieves the best performance with the minimum number of trainable parameters (0.237 million) and runtime (0.0023s per image). Additionally, larger models impose higher requirements on both training and testing, demanding more powerful and expensive hardware, which can hinder their wide applicability.

**Table 1.** The Runtime/parameters comparison between models. As the number of parameters (in millions) increases, the runtime (in seconds) also increases. DeBCR achieves better performance in both metrics with 0.237 million parameters and a runtime of 0.0023 seconds. Notably, DDPM ranks at the bottom in terms of runtime because this model relies on the diffusion process, which consumes significant time.

| Model | DeBCR | CARE | DnCNN | N2N | UniFMIR | U-Net | RCAN | MPRNet | DDPM | ESRGAN | TA-GAN |
| --- | --- | --- | --- | --- | --- | --- | --- | --- | --- | --- | --- |
| Params. | **0.237** | 0.333 | 0.556 | 1.227 | 1.565 | 7.780 | 15.334 | 20.127 | 23.998 | 49.841 | 30.782 |
| Runtime | **0.0023** | 0.0039 | 0.0056 | 0.0068 | 0.0042 | 0.0241 | 0.0281 | 0.0173 | 1.0968 | 0.0147 | 0.0067 |

## Supplementary material

## Comparison between the vanilla structure of BCR to residual BCR

A BCR decomposition unit at certain decomposition levels can be represented with the DNN structure in Fig.S1a below. The LC indicates the local convolution and the conv2D is the normal 2D convolution in the DNN toolset (Tensorflow in this work). Yet, this vanilla structure in Fig.S1a shows inferior stability during the restoration process. The reason behind this lies in the inherent nonlinearity of the microscopy imaging system in reality, whereas the BCR theory necessitates linearity in the question configurations. To enhance the non-linear properties of the network, we introduced the residual BCR decomposition unit as Fig.S1b.

**Fig. S1.**
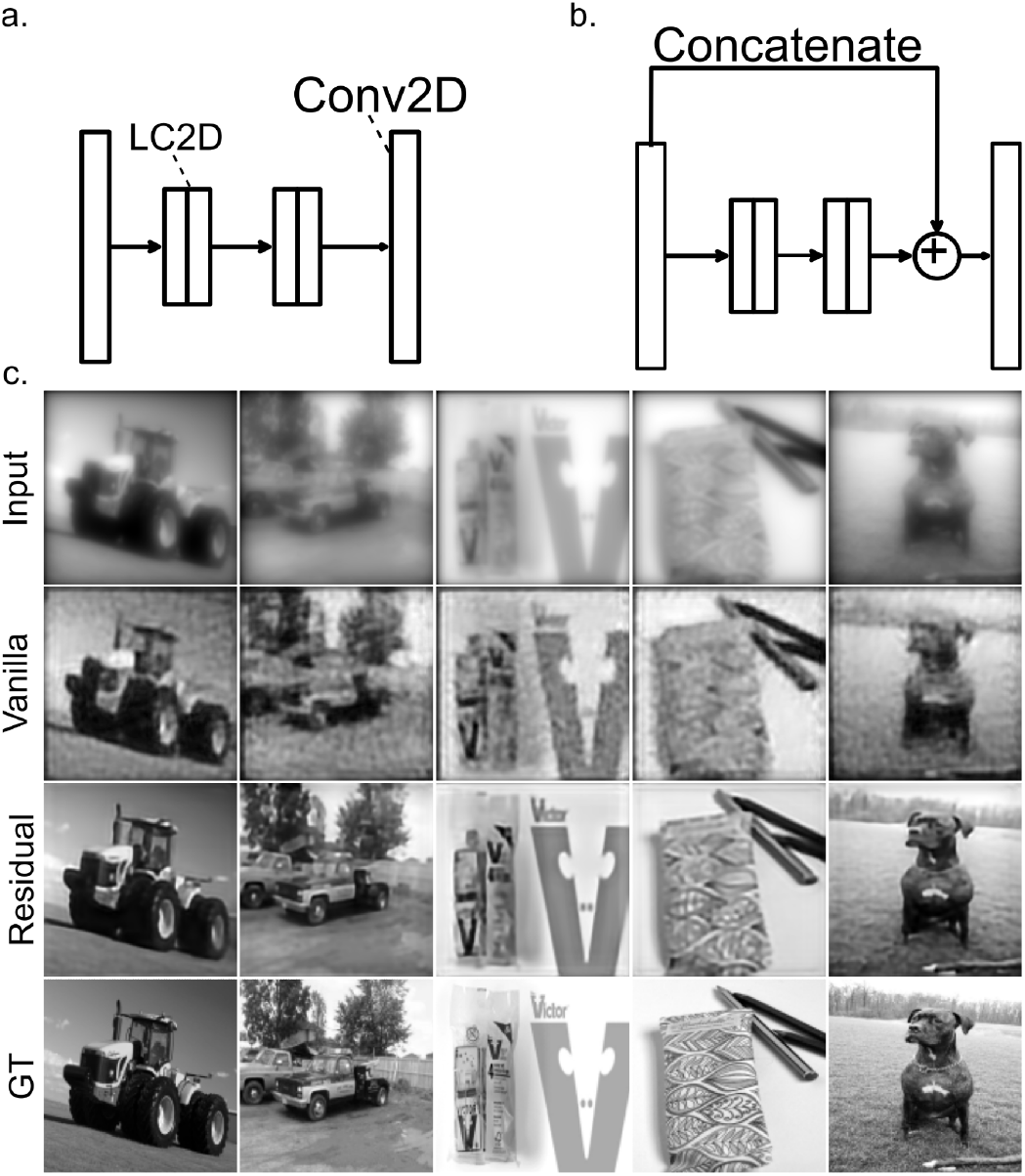
The BCR decomposition unit for the integral operator. **a**, the original BCR decomposition unit. LC2D indicated the 2D local convolution. The approximation based on this structure requires the target system to be linear. **b**, The residual BCR unit for non-linear problems. The microscopy imaging system inherently conforms to non-linear properties due to the prevalence of dominant disturbing elements ε. Incorporating the residual structure assists in rendering the task more robust. **c**, the comparison between vanilla BCR unit and residual BCR by image restoration on ImageNet. The vanilla BCR unit is less effective at compressing artifacts during reconstruction, whereas the residual BCR unit stabilizes the learning process.

### Network architecture

Building upon the preceding DNN structure in Fig.S1b for the BCR decomposition unit, we can represent the inverse operator in integral operation form in Equation (5) with the DeBCR model in Fig.S.2. The input images undergo two rounds of downsampling for multi-stage learning. At each stage, the image undergoes BCR-based decomposition equations. The resulting product yields the restored image at the corresponding resolution level. In contrast to the traditional single-stage residual BCR learning process, DeBCR integrates feature maps from other resolution levels. Leveraging posterior information from these additional resolution levels, the model effectively stabilizes the learning process.

**Fig. S2.**
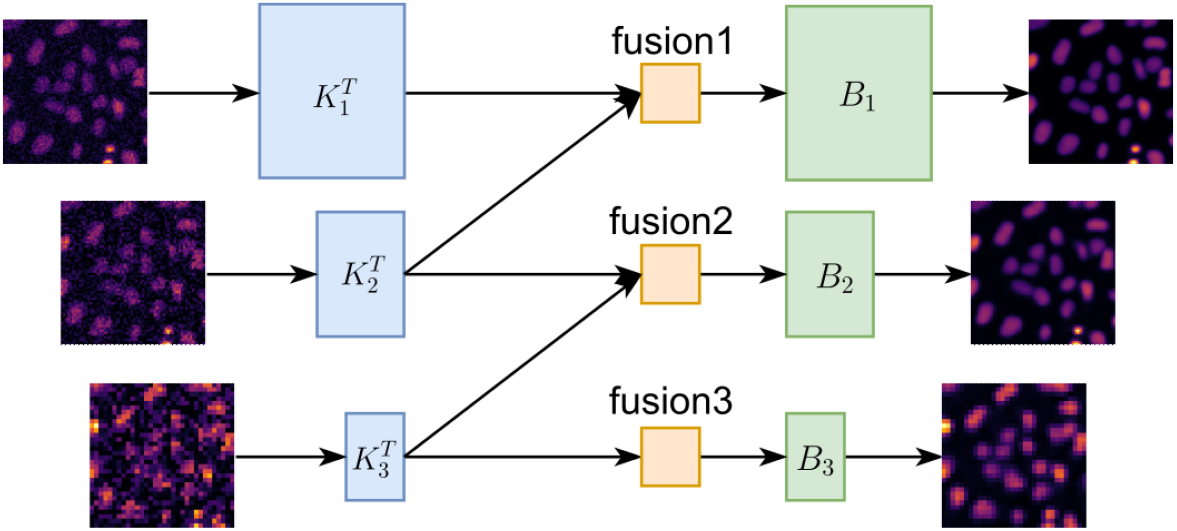
Multi-stage decomposition of the DeBCR models. The DeBCR contains multiple stages of decomposition of the restorations. The noisy input will be subjected to two rounds of downsampling to serve as inputs for the other two resolution levels. All reconstructions adhere to the previous decomposition conclusions outlined in equation (5). 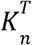 indicates the integral operator in *n* downsampling resolution. The *B*_*n*_ corresponds to its pseudo-differential operator. In addition to the single decomposition, the model incorporates feature maps from other resolution levels to stabilize learning performance through the fusion operator contain the restorations in multiple resolutions. *Fusion*_*n*_. The outputs contain the restorations in multiple resolutions.

### Comparison to the newly-released benchmark UniFMIR in LM

We compared the DeBCR to the newly released benchmark for the microscopy restoration model UniFMIR[1]. UniFMIR achieved restoration through pre-training on a large base model, which necessitated significant computational resources. Training the model from scratch for comparison purposes would be challenging. Therefore, our comparison of UniFMIR’s performance relies solely on the pre-trained weights provided by the author. The UniFMIR models encompass two overlapping experiments to DeBCR. These include the flatworm *Schmidtea mediterranea* datasets for brightfield microscopy 2D denoising and the *F-Actin* dataset for confocal microscopy super-resolution (SR). Results are presented in Fig.S3.

Under medium/weak signal-to-noise ratio (SNR) conditions (C1, C2), even though the restorations differ from each other, both DeBCR and UniFMIR effectively restored information from noisy inputs (Fig.S3a). However, as the SNR decreases to the extremely weak condition (C3), UniFMIR exhibits tendencies toward restoring the fake patterns as indicated by the green arrow. In cases where the input consists solely of noise (fourth column), where no signals exist in the ground truth (GT), UniFMIR still generates artifacts due to hallucinations. The DeBCR correctly identifies that there is only noise in the inputs. The inferior capacity to suppress hallucinations results in poorer evaluation scores for UniFMIR compared to DeBCR. Specifically, in terms of PSNR↑/SSIM↑ evaluation metrics, DeBCR outperforms UniFMIR with scores of 29.04/0.92 compared to 24.36/0.72. Both models demonstrate strength in restoring high-resolution details from low-resolution inputs. However, DeBCR exhibits superior artifact constraint and restores images closer to the ground truth (GT). Evaluation metrics such as PSNR↑/SSIM↑ further validate DeBCR’s performance, with scores of 27.01/0.84, surpassing UniFMIR’s scores of 25.74/0.77.

In the SR task on the dataset confocal and the stimulated emission depletion microscopy (STED) for *F-Actin* (Fig.S3b) Both models demonstrate strength in restoring high-resolution details from low-resolution inputs. However, DeBCR exhibits superior artifact constraints and restores images closer to the ground truth (GT). Evaluation metrics such as PSNR↑/SSIM↑ further validate DeBCR’s performance, with scores of 27.01/0.84, surpassing UniFMIR’s scores of 25.74/0.77.

**Fig. S3.**
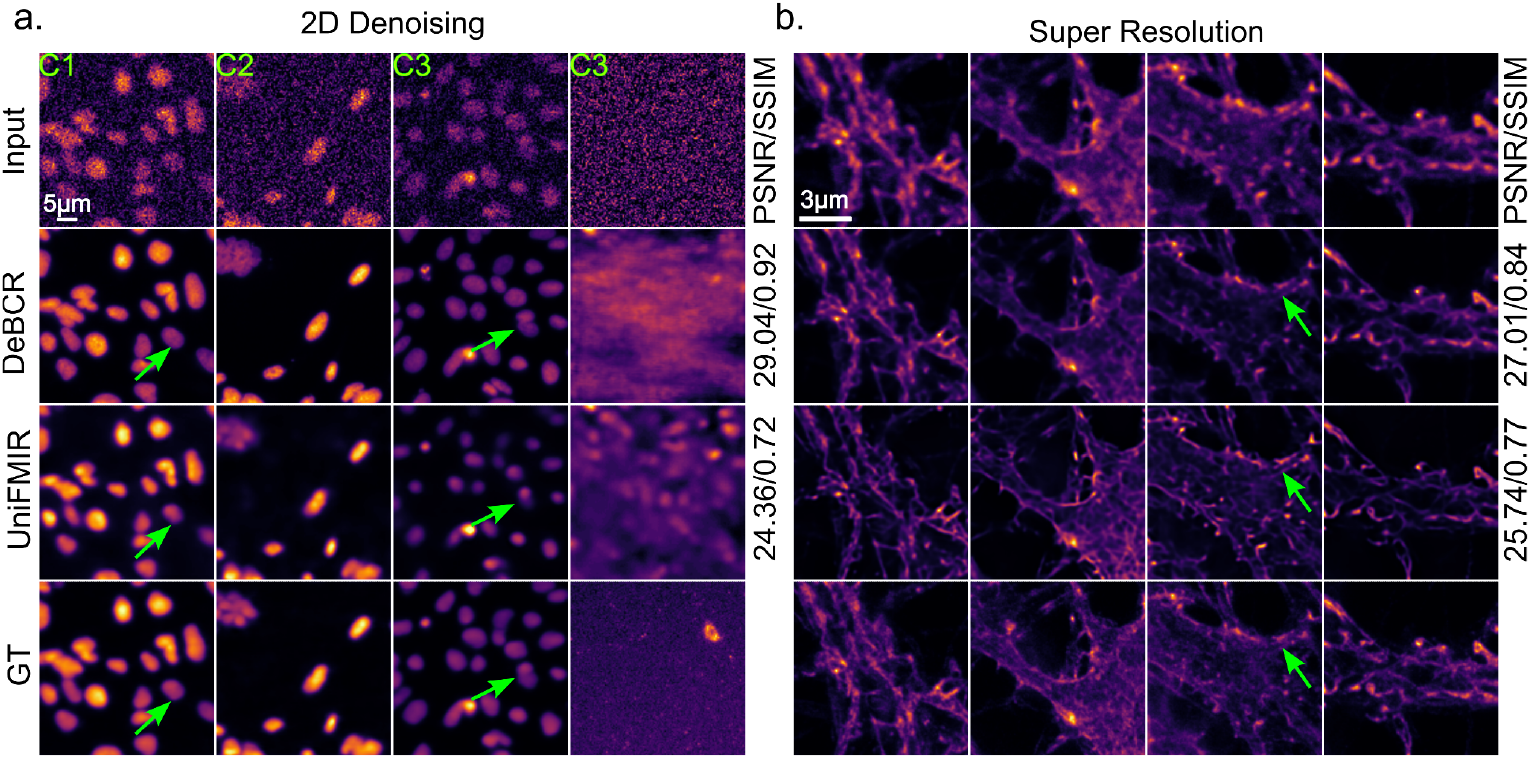
Deconvolution and denoising performance compared to the universal fluorescence microscopy-based image restoration (UniFMIR[1]). **a**, denoising results on the flatworm *Schmidtea mediterranea* datasets. **b**, the SR task on the dataset confocal and the stimulated emission depletion microscopy (STED) for *F-Actin*.

### Spectral effect of DeBCR restoration observed on cryo-ET tomograms data

As was described before, the spectral noise distribution can be estimated from same-signal noise-independent even/odd volumes by obtaining the difference between volume V_even_-V_odd_, while the sum volume V_even_+V_odd_ would serve as the full (i.e. noisy signal) volume of the same noise level[2]. Then, the radially averaged power spectra for full and estimated noise are obtained and normalized to the corresponding full power spectrum energy.

The DeBCR-denoised data exhibits nonlinear frequency response on DeBCR filtering with the decreased estimated noise and the increased full content power in the low- and mid-frequency ranges (Fig.S4). At the same time, DeBCR acts as a low-pass filter with a smooth transition band (Fig.S4) at the higher frequencies.

**Fig. S4.**
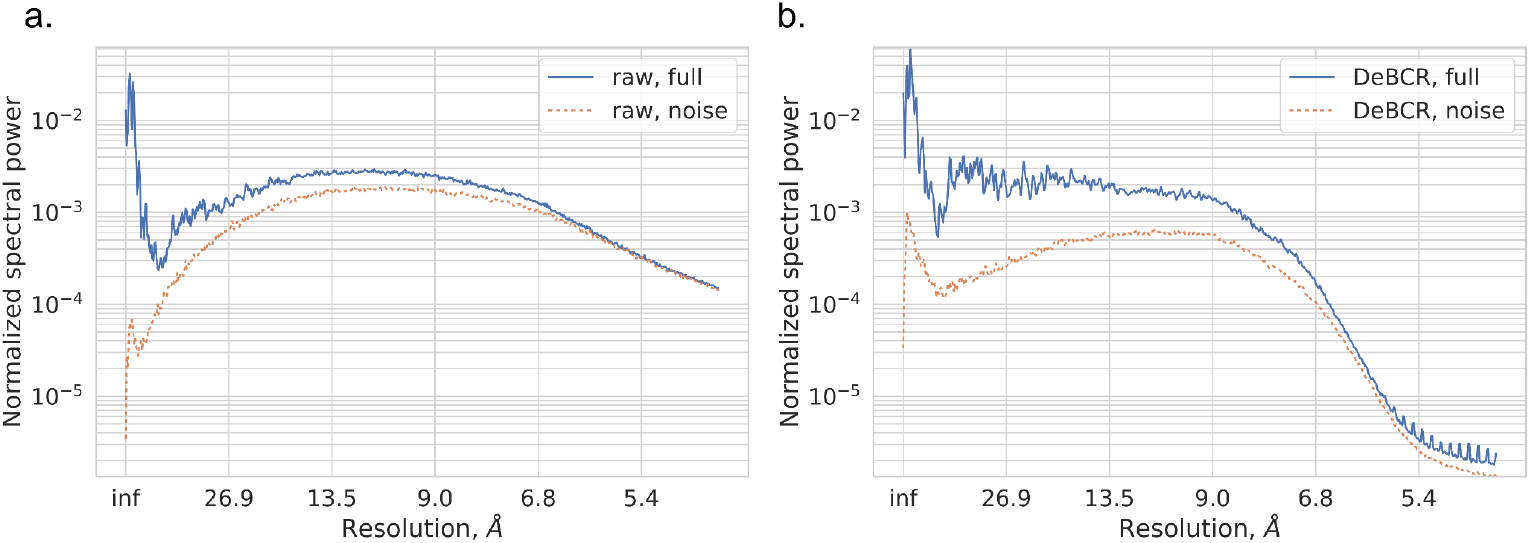
Spectral comparison of raw and DeBCR-denoised cryoET data. (Tomo110 dataset, Fig.5a-b). **a**, Normalized radially averaged full and estimated noise power spectra for the raw data; **b**, The DeBCR-denoised data.

### Subtomogram averaging of RyR1 particles from raw and DeBCR denoised data

Even- and odd frames from the tilt series were processed separately by tomoBEAR[3]. In brief, frames were motion-corrected by MotionCor2[4], tilt series were aligned by IMOD[5], and defocus values were detected by Gctf[6]. Even and odd half tomograms were generated by weighted back-projection in IMOD[5]. Locations o

The RyR1 molecules were detected by Dynamo[7]-style template matching implemented in tomoBEAR[3] in CTF-phase-flipped tomograms. Furthermore, Dynamo multi-reference alignments were applied to exclude the false positive segments from template matching and align all the RyR1 particles to the same orientation. Finally, 34 particles were manually selected from 22 even-odd pairs of tomograms for the FSC analysis (Fig.6e).

The same process of subtomogram averaging was implemented on DeBCR denoised tomograms and raw tomograms. Benefiting from the signal enhancement of DeBCR, the 3D subtomogram average reconstruction from denoised data demonstrated better visible contrast (Fig.6d) while preserving the high-resolution information compared to the classical linear bandpass filtering (Fig.6e). With the same EM volume intensity threshold values and quite limited amount of particles, the DeBCR-denoised structure exhibits more molecular density features than the original data in Fig.S5. This result bears a promising potential of reducing requirements on amounts of particles to achieve certain resolution and, subsequently, amounts of the collected and processed data. Therefore, DeBCR-denoised tomograms could be used to generate initial subtomogram averages, however, high-resolution refinement should be done on raw data.

**Fig. S5.**
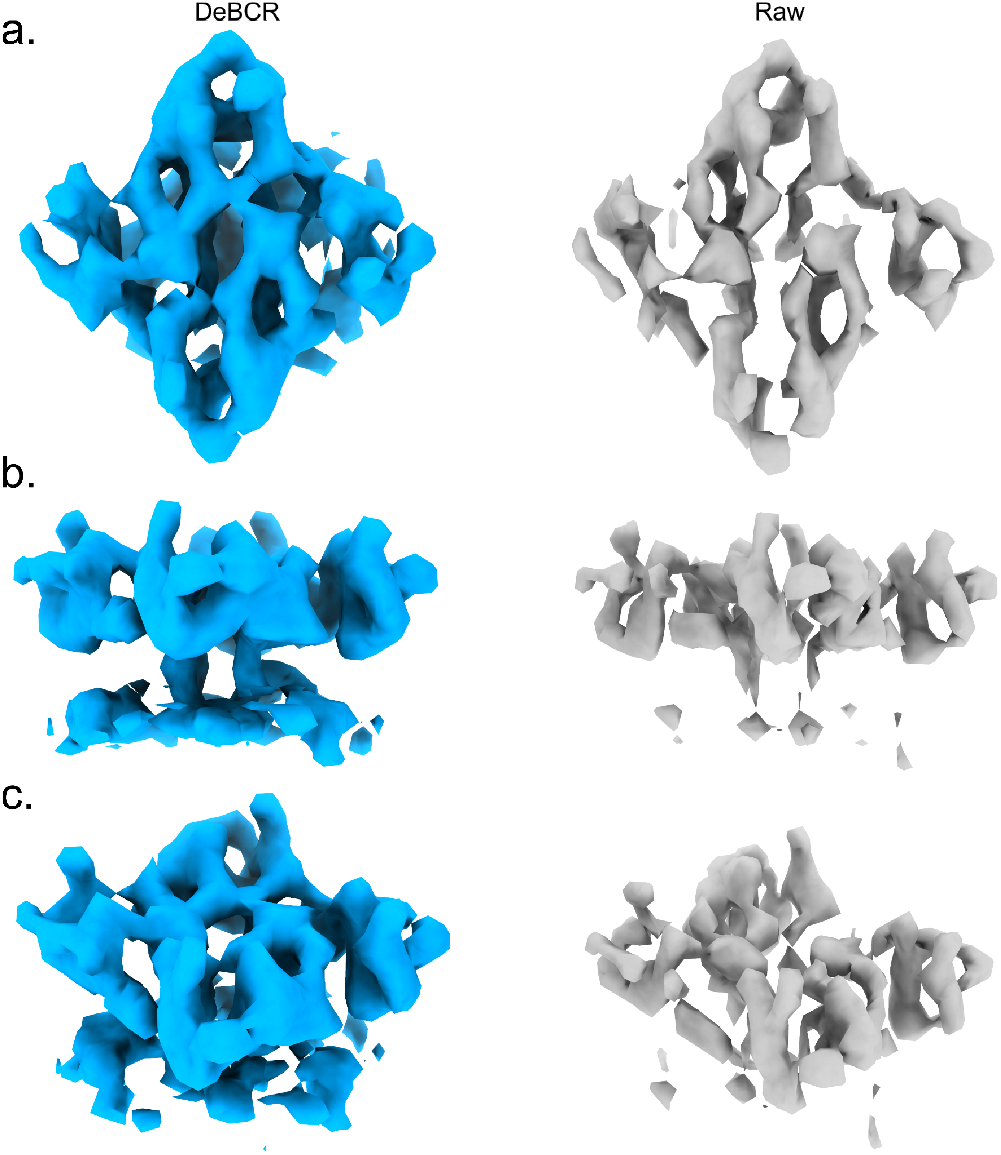
The 3D molecular map of RyR1 reconstruction by averaging 34 manually selected particles obtained from the raw and DeBCR-restored tomograms for the RyR1 in native vesicles dataset (EMPIAR-10452). The averaging results were thresholded to the same value in ChimeraX. **a**, the top view. **b**, the side view. **c**, the oblique view.

